# Sensory computations in the cuneate nucleus of macaques

**DOI:** 10.1101/2021.07.28.454185

**Authors:** Aneesha K. Suresh, Charles M. Greenspon, Qinpu He, Joshua M. Rosenow, Lee E. Miller, Sliman J. Bensmaia

## Abstract

Tactile nerve fibers fall into a few classes that can be readily distinguished based on their spatiotemporal response properties. Because nerve fibers reflect local skin deformations, they individually carry ambiguous signals about object features. In contrast, cortical neurons exhibit heterogeneous response properties that reflect computations applied to convergent input from multiple classes of afferents, which confer to them a selectivity for behaviorally relevant features of objects. The conventional view is that these complex response properties arise within the cortex itself, implying that sensory signals are not processed to any significant extent in the two intervening structures – the cuneate nucleus (CN) and the thalamus. To test this hypothesis, we recorded the responses evoked in CN to a battery of stimuli that have been extensively used to characterize tactile coding in both the periphery and cortex, including skin indentations, vibrations, random dot patterns, and scanned edges. We found that CN responses are more similar to their cortical counterparts than they are to their inputs: CN neurons receive input from multiple classes of nerve fibers, they have spatially complex receptive fields, and they exhibit selectivity for object features. Contrary to consensus, then, CN plays a key role in processing tactile information.

**Significance:** Perception is the outcome of the sequential processing of sensory signals at multiple stages along the neuraxis. The conventional view is that tactile signals are processed predominantly in the cerebral cortex. We tested this view by investigating the response properties of neurons in the cuneate nucleus (CN), the first potential stage of processing along the primary touch neuraxis. We found that CN responses more nearly resemble those of cortical neurons than they do those of nerve fibers: CN neurons have spatially complex receptive fields reflecting convergent input from multiple classes of nerve fibers and exhibit a selectivity for object features, absent in the nerve. We conclude that CN plays a key, early role in the processing of tactile information.

## Introduction

The coding of tactile information has been extensively studied in the peripheral nerves and in the primary somatosensory cortex (S1, Broadmann’s area 3b) of non-human primates, leading to the conclusion that sensory representations in S1 differ from those at the periphery in at least two important ways. First, while cutaneous nerve fibers can be divided into a small number of classes, each responding to a different aspect of skin deformation, individual S1 neurons integrate sensory signals from multiple classes of nerve fibers (Lieber and Bensmaia, 2019; Pei et al., 2009; Saal and Bensmaia, 2014). Indeed, while each class of nerve fibers exhibits stereotyped responses to certain stimulus classes, for example skin indentations or sinusoidal vibrations, cortical responses to these same stimuli include features of the responses from multiple tactile classes, or sub-modalities. Second, the responses of cortical neurons reflect computations on these inputs. For example, the spatial receptive fields of S1 neurons comprise excitatory and inhibitory subfields, implying a spatial computation (Bensmaia et al., 2008; DiCarlo et al., 1998). Similarly, S1 neurons act as temporal filters, as evidenced by the fact that their responses to vibrations reflect both integration and differentiation of their inputs in time (Saal et al., 2015). These computations give rise to increasingly explicit rate-based representations of object features, such as the orientation of an edge indented into the skin or the texture of a surface scanned across the skin (Bensmaia et al., 2008; Lieber and Bensmaia, 2019).

In contrast to the well-studied peripheral and cortical representations of touch, very little is known about the contribution of the cuneate nucleus (CN) to the processing of tactile information. The textbook view is that CN acts as a simple relay station despite the fact that the response properties of neurons in CN or equivalent brain structures (nucleus principalis, e.g.) exhibit responses that are not identical to those of nerve fibers (Bystrzycka et al., 1977; Ebert et al., 2021; Jörntell et al., 2014; Kaloti et al., 2016; Witham and Baker, 2011), implying some processing. However, CN responses have not been investigated using stimuli whose representation in the nerve and cortex has been quantitatively characterized (Conner et al., 2021; Ebert et al., 2021; Jörntell et al., 2014; Lehnert et al., 2021). This precludes a quantitative analysis of how tactile signals are transformed in this structure.

To fill this gap, we recorded the responses evoked in individual CN neurons to a battery of tactile stimuli that have been extensively used to characterize the response properties of tactile nerve fibers and of neurons in S1, including skin indentations, vibrations, embossed dot patterns, and scanned edges. We then compared CN responses to their upstream (nerve fibers) and downstream counterparts (area 3b or S1, the first stage of processing in cortex (Delhaye et al., 2018; Kaas, 1983)) to assess the degree to which tactile signals are processed in CN. The picture that emerges is one in which CN plays an integral part in the transformation of tactile information as it ascends the neuraxis.

## Results

To investigate tactile representations in CN, we measured the responses of individual CN neurons (n=33) to step indentations, sinusoidal skin vibrations (n=68), mechanical noise (n=33), random dot patterns (n=31), and scanned bars (n=9). To compare CN responses to their peripheral counterparts, we simulated the spiking responses of tactile nerve fibers to the stimuli used in the CN recordings using a model that can reconstruct such responses with millisecond-level precision (Saal et al., 2017). To compare CN responses to their cortical counterparts, we analyzed previously collected cortical responses to analogous stimuli.

### Adaptation properties of CN neurons reveal submodality convergence

Nerve fibers can be readily divided into two groups based on their responses to skin indentations: Slowly adapting type 1 (SA1) fibers respond throughout the skin indentation whereas rapidly adapting (RA) and Pacinian Corpuscle-associated (PC) fibers respond only to the onset and offset of the indentation and are silent during the intermediate sustained epoch (Pei et al., 2009). Examination of the responses of downstream neurons to skin indentations can thus reveal the sub-modality composition of their inputs. Specifically, responses during the sustained component reflect SA1 input, as only this class is active during this stimulus epoch; a strong phasic response during the offset of the indentation is indicative of RA or PC input, as only these two classes of nerve fibers produce an off response. Co-occurrence of these two response properties reflects convergent input from at least two classes of nerve fibers. In CN, we found that the responses of a majority of neurons comprise both sustained and off components, indicative of convergent input from multiple sub-modalities (Figure 1a).

**Figure 1.**
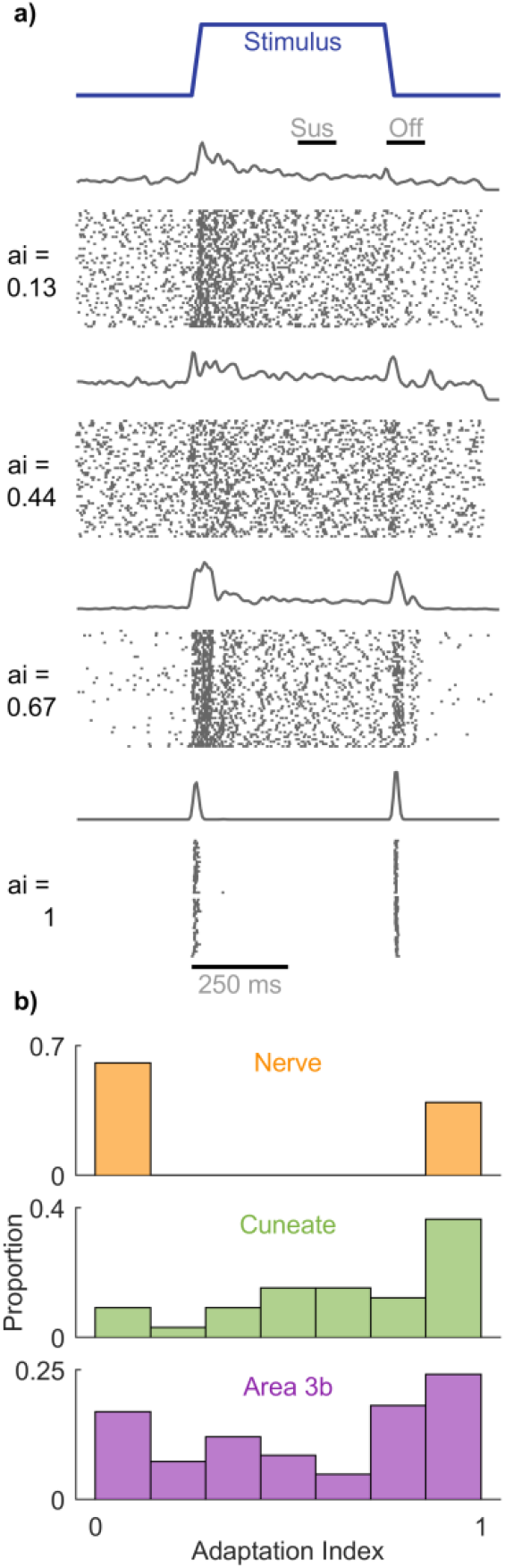
CN responses to step indentations. a| Responses of four CN neurons that span the range of convergence properties. b| AI index for the nerve (top), cuneate nucleus (middle), and the primary somatosensory cortex (bottom). AI segregates nerve fibers at the two extremes, whereas convergence is observed in both the CN and S1.

A previously developed ‘adaptation index’ (Pei et al., 2009) gauges the degree to which individual neurons receive convergent input from multiple cutaneous sub-modalities based on the relative strengths of the sustained and off responses. A value of 1 denotes RA-like responses (only an off-response, no sustained response), a value of 0 denotes SA1-like responses (only a sustained response, no off response), and an intermediate value denotes convergent input (mixture of sustained and off responses). Adaptation indices computed on CN responses spanned the range from 0 to 1, with most falling between the two extremes, suggesting that convergence is the rule rather than the exception (Figure 1b). A greater number of neurons exhibited pure RA-like than SA1-like responses, as has been found in S1, commensurate with the relative densities of these two groups of nerve fibers (RA/PC vs. SA1). To obtain a quantitative estimate of the proportion of multimodal neurons, we tested whether the firing rates during the sustained and offset periods were significantly different from the baseline period. Of the 33 neurons tested with skin indentations, 6% produced only sustained responses, 27% only offset responses, and 60% produced both sustained and offset responses (the remaining 7% only produced a transient onset response). Convergence of cutaneous sub-modalities is thus observed in a majority of neurons in CN.

### CN responses to vibrations reveal submodality convergence

Next, we examined the responses of CN neurons to sinusoidal vibrations varying in amplitude and frequency, hoping to capitalize on the fact that different afferent classes exhibit different frequency sensitivity: SA1 fibers peak in sensitivity at low frequencies, PC fibers at high frequencies, and RA fibers at intermediate frequencies (Muniak et al., 2007; Talbot et al., 1968). We can then assess whether the frequency response characteristic of individual CN neurons resembles that of any single class of tactile nerve fiber or rather reflects convergent input from multiple fiber types. We found that some CN neurons respond exclusively to low frequencies, similar to SA1 fibers (Figure 2a), others to high frequencies, similar to PC fibers (Figure 2b), but many respond to the entire range of frequencies tested (Figure 2c), suggesting they receive convergent input from multiple tactile sub-modalities.

**Figure 2.**
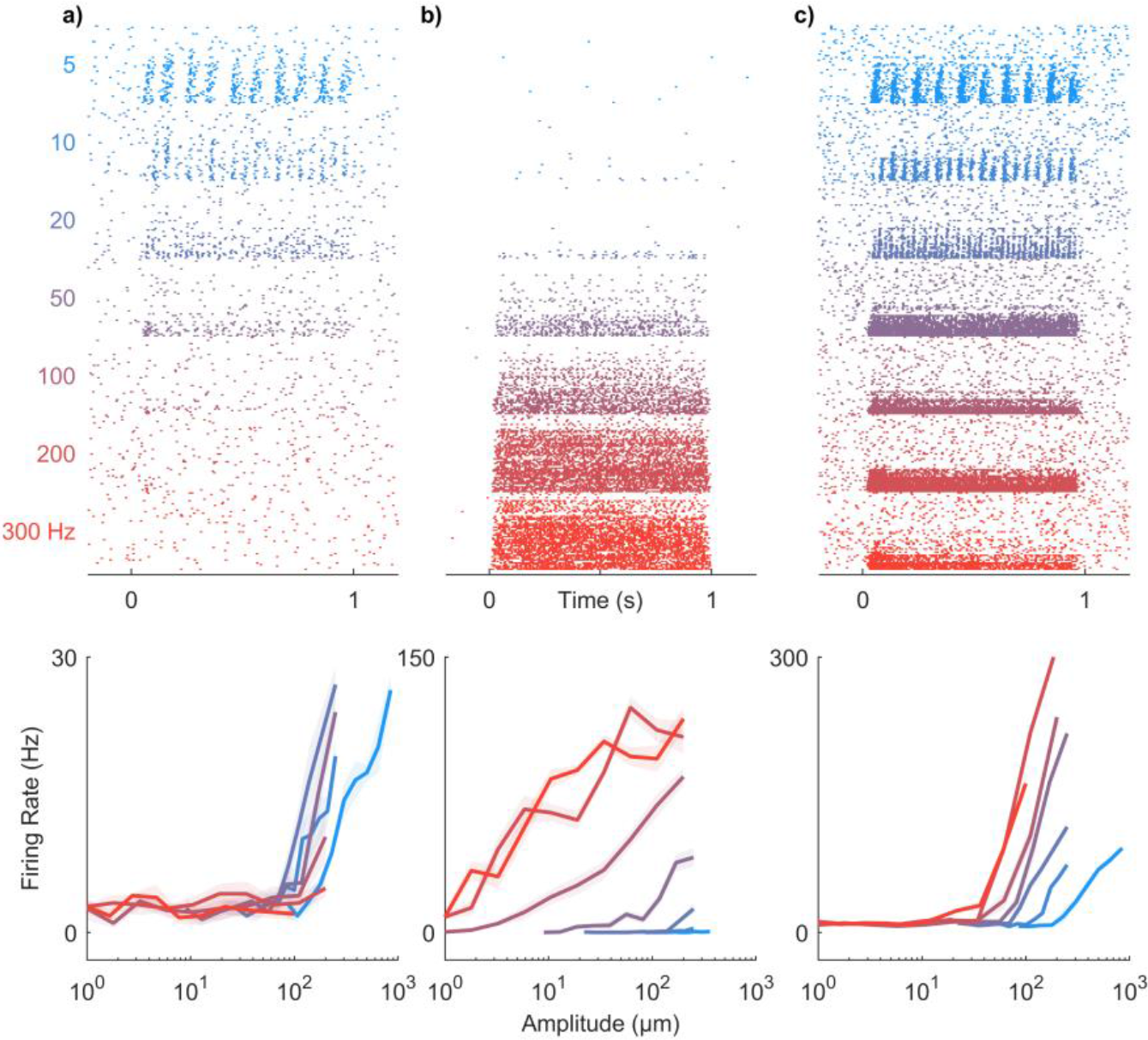
CN responses to vibrations. a-c| Responses of three CN neurons to skin vibrations varying in frequency (from 5 to 300 Hz) and amplitude (1-1000 µm; ordered by frequency, then amplitude). Some CN neurons responded exclusively at low frequencies (a), others at high frequencies (b), but many CN neurons responded over a wider range of frequencies than does any one population of nerve fibers (c). As is the case in periphery and cortex, CN neurons often exhibited phase-locked responses to vibratory stimuli (see Supplemental Figure 1).

To quantitatively assess the contributions of different afferent classes to the responses of CN neurons, we regressed the firing rates of individual CN neurons onto the (simulated) population firing rates of nerve fibers from all three classes to a common set of vibrations (Figure 3a). First, we verified that the responses of most CN neurons could be well accounted for using a linear combination of SA1, RA, and PC responses (mean R^*2*^=0.6). Second, we assessed whether CN responses were better accounted for by multiple afferent classes than by one and found that, for most CN neurons, the cross-validated model fit increased significantly with the inclusion of all inputs (Figure 3a, mean R^2^_best_ = 0.50, mean R^2^_all_ = 0.60, mean ΔR^2^ = 0.1, ranksum = 5395, z = 3.2, p = 0.0013). We repeated the regression analysis on measured responses of tactile nerve fibers to similar sinusoidal vibrations to verify our ability to distinguish unimodal from multimodal responses. We found that measured afferent responses to vibrations were equally well accounted for with a single modality as they were multiple modalities (Figure 3a, mean R^2^_best_ = 0.83, mean R^2^_all_ = 0.86, mean ΔR^2^ = 0.03, ranksum = 1019, z = 1.53, p = 0.126). We found that 46% of CN neurons yielded ΔR^2^ that were more than one standard deviation away from the mean ΔR^2^ obtained from nerve fibers, whereas only 10% of nerve fibers exceeded this threshold.

**Figure 3.**
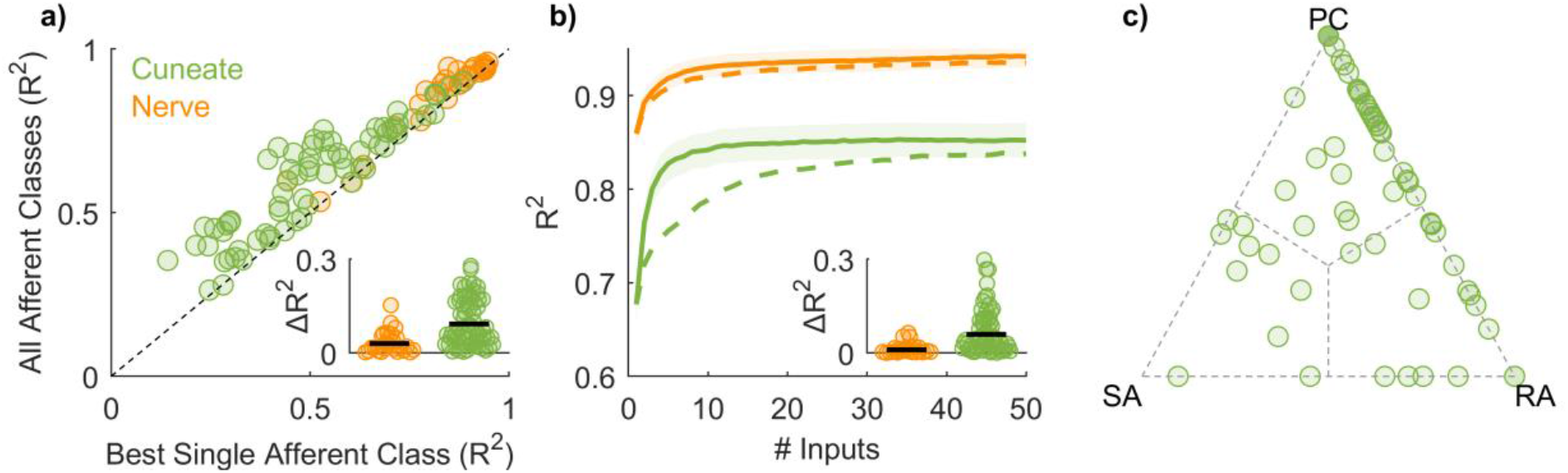
CN responses to vibrations reflect convergent input from multiple afferents, typically of multiple classes. **a**| Model fit with only one class of nerve fibers vs. model fit with multiple classes for CN and afferents. The responses of nerve fibers can be predicted nearly perfectly with a single afferent type whereas CN neurons often require multiple. Inset: model improvement when allowing all classes is significantly greater for cuneate than individual afferents. For this analysis, the mean response to each stimulus is used as a regressor, computed separately for each class of nerve fibers. b| Performance of regression models as a function of the number of afferents included in the analysis. Input from 2-5 nerve fibers is sufficient to achieve asymptotic performance for CN predictions, but only if convergence across sub-modalities is allowed. When only the best single afferent class is used (dashed line), an order of magnitude more afferents are required to reach asymptotic performance. Inset: At criterion, model performance is significantly improved when all afferent classes are included as regressors in models of CN responses. c| Normalized regression weights for each afferent class; each point corresponds to a CN neuron.

Third, we estimated the number of afferent inputs required to predict CN responses accurately. To this end, we simulated the responses of a population of nerve fibers and assessed our ability to predict the responses of individual CN neurons as we sequentially added simulated nerve fibers to the regression model (Figure 3b). We found that model fits typically leveled off (reached criterion performance) with just 2-5 inputs if all three classes of nerve fibers were included in the analysis. If only the most predictive afferent class was included, more inputs were required to achieve equivalent fits and performance plateaued at a lower level, consistent with the above analysis based on mean (simulated) population responses (Figure 3b – dashed line). Including all three afferent classes as regressors significantly improved CN predictions (mean ΔR^2^ = 0.06, ranksum = 5518, z = 3.07, p = 0.002). We validated the approach by verifying that including all three classes did not improve afferent predictions (mean ΔR^2^ = 0.01, ranksum = 977, z = 0.90, p = 0.36). Examination of the optimized regression coefficients revealed that 15% of CN neurons were unimodal, 59% were bimodal, and the remainder (26%) were trimodal (Figure 3c). In conclusion, then, the responses of individual CN neurons to vibrations reflect input from multiple classes of nerve fibers so the submodality convergence observed in cortex is at least in part inherited from CN.

### CN responses to vibrations reveal temporal computations

Neurons in somatosensory cortex have been shown to exhibit a variety of response properties to vibrations (Saal et al., 2015). Some neurons sum their inputs over time whereas others act as more complex temporal filters, comprising both excitatory and suppressive components. Examination of the rate-intensity functions for vibrations revealed suppressive components in the neuronal response (Supplementary Figure 1): Some CN neurons were always suppressed by vibration whereas others were excited by some vibrations and suppressed by others. For these neurons, regression models yielded significantly poorer fits when weights were constrained to be positive (mean 22% decrease). These suppressive components may constitute building blocks of more complex temporal feature filtering.

To further characterize the process of temporal integration, we examined CN responses to mechanical noise. Specifically, we computed the mean response evoked in each afferent class immediately preceding each spike evoked in a given CN neuron (Figure 4). The resulting spike-triggered averages represent how a neuron integrates the signal from each population of nerve fibers. Some neurons simply summed their afferent input whereas others exhibited more complex response properties, with STAs that comprised excitatory and suppressive components, similar to those derived from S1 responses to analogous stimuli. As is the case in cortex, PC input tended to be more suppressive than was RA or SA1 input (Figure 4c). Temporal receptive fields that comprise excitatory and suppressive components confer to neurons a preference to fluctuations in the afferent input; heterogeneity in the filters across neurons and input classes (cf. (Saal et al., 2015)) gives rise to a high-dimensional representation of the input (Lieber and Bensmaia, 2019).

**Figure 4.**
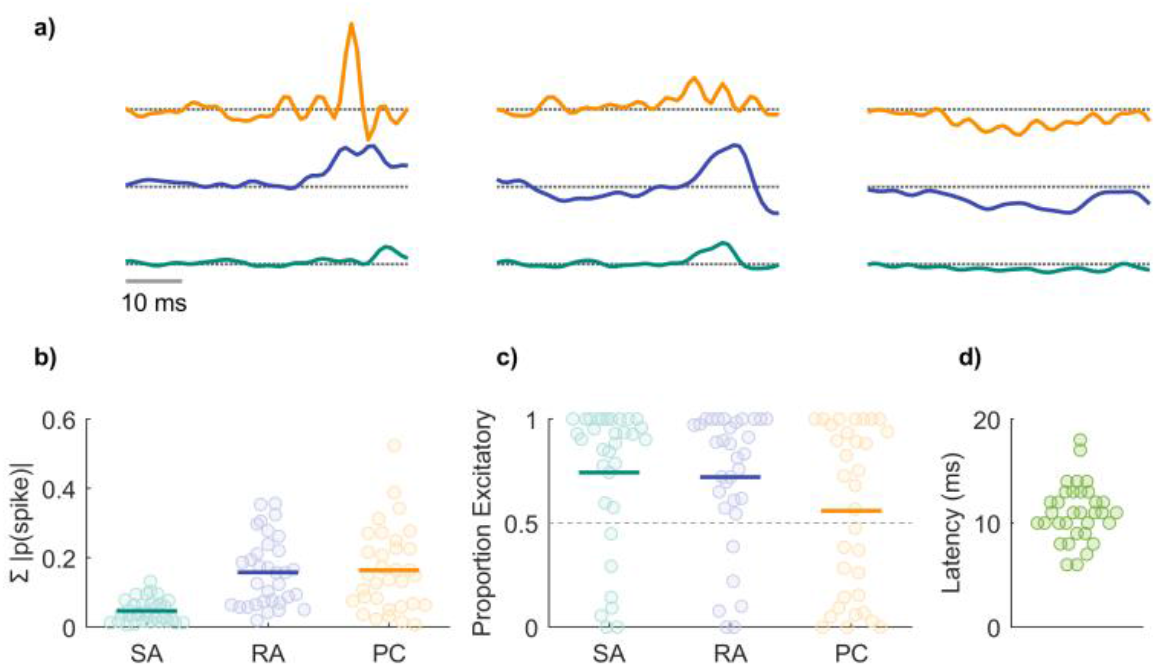
Temporal integration properties of CN neurons. a| Spike-triggered averages (STAs) computed from the responses of 3 CN neurons for inputs from the three classes of nerve fibers. STAs comprise both excitatory and suppressive components, as do their counterparts derived from the responses of S1 neurons. b| Summed absolute spike probability for each CN neuron with respect to afferent type. Given the frequency composition of the vibrations, the RA and PC drive was greater than was SA1 drive. c| Proportion of the afferent input that is excitatory vs suppressive. The temporal receptive fields of many CN neurons included both excitatory and suppressive components d| The latency, estimated from responses to mechanical noise, was about half of that in S1.

From CN responses to mechanical noise, we also estimated the mean latency in CN to be around 10 ms (Figure 4d), approximately half of that in S1 (∼18 ms).

### Spatial structure of CN receptive fields

Neurons in somatosensory cortex act not only as temporal filters (Saal et al., 2015) but also as spatial filters (Bensmaia et al., 2008; DiCarlo et al., 1998; Lieber and Bensmaia, 2019). The spatial receptive fields of S1 neurons comprise excitatory and inhibitory subfields, conferring to them a sensitivity to specific spatial features in their inputs. For example, an elongated excitatory subfield flanked by an inhibitory one will confer to a neuron a selectivity for orientation (Bensmaia et al., 2008; Hubel and Wiesel, 1962).

With this in mind, we reconstructed the spatial receptive fields of CN neurons from their responses to random patterns of embossed dots scanned across the skin (cf. (DiCarlo et al., 1998; Lieber and Bensmaia, 2019)). First, we found that the RFs of CN neurons tend to be larger than are those of SA1 or RA fibers (Johnson and Lamb, 1981), as expected given the inferred convergence of afferent input onto individual CN neurons (Figure 5a,b). Second, CN neurons have marginally smaller RFs than do their cortical counterparts (Figure 5b,d), as expected given their relative positions along the neuraxis. The mean RF size is 7.2 mm^2^ in CN and 9.9 mm^2^ in cortex (t-test: t(44) = 2.8, p = 0.072). Third, CN neurons have complex RFs, often comprising excitatory and inhibitory subfields, like their cortical counterparts (Figure 5a,c). As in cortex, the excitatory subfields of CN neurons tend to be larger than their inhibitory counterparts. However, CN RFs tend to comprise a greater number of distinct subfields than do their S1 RFs (mean of 4 vs. 2.1 mm^2^, t(44) = 4, p < 0.001). While the excitatory masses are similar in CN and S1, the inhibitory masses are smaller in CN than in S1 (excitatory: 5.6 vs. 6.6 mm^2^ t(44) = 1.37, p = 0.178; inhibitory: 1.5 vs. 3.3 mm^2^, t(44) = 2.8, p < 0.008). Nonetheless, the spatial structure of the receptive fields observed in CN is qualitatively similar to its counterpart in S1.

**Figure 5.**
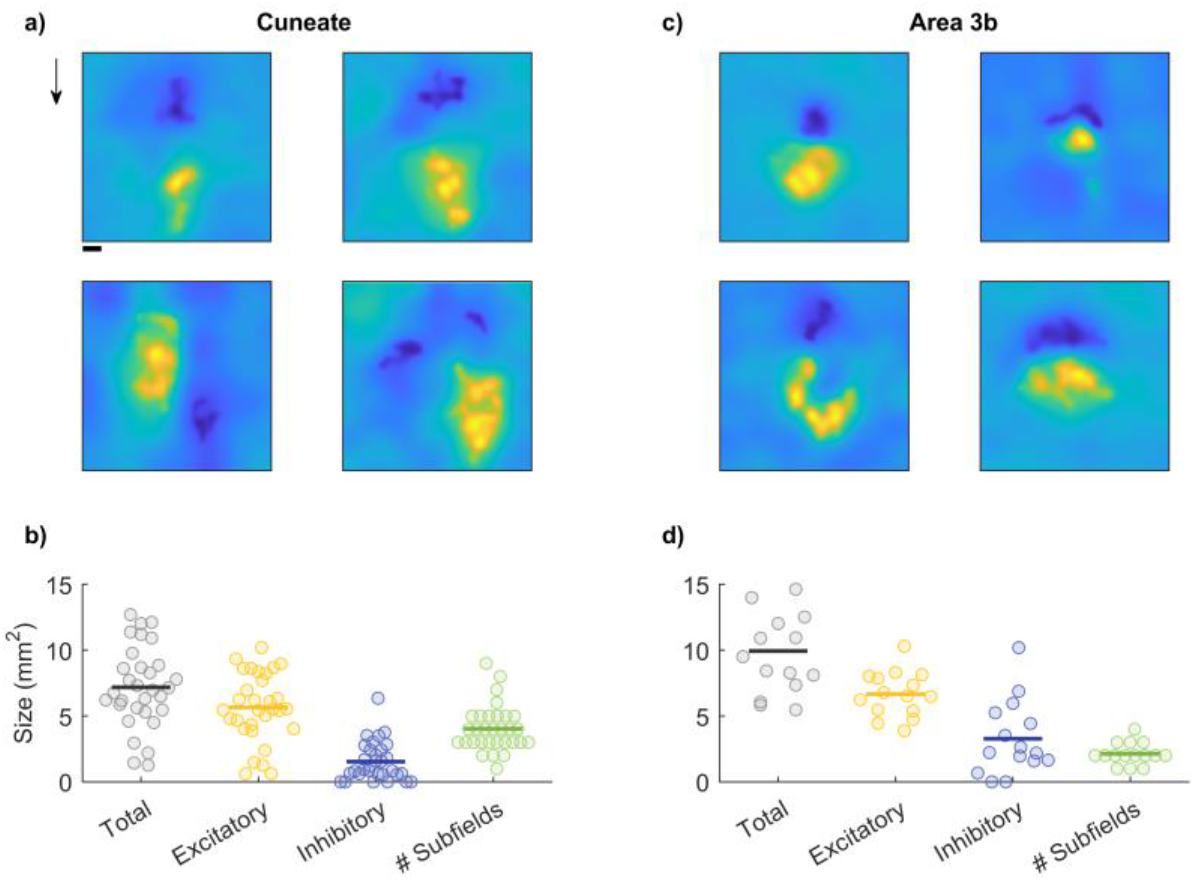
Spatial receptive fields of CN neurons. a| Reconstructed receptive fields for 4 cuneate neurons. RFs typically comprise both excitatory and inhibitory subfields in a variety of conformations. Cuneate RFs are similar to their S1 counterparts (panel c). Arrow indicates the direction in which the dot pattern was scanned. Scale bar is 1mm. b| Cuneate receptive fields are on average smaller than those in S1 (panel d), a difference that is primarily driven by smaller inhibitory subfields.

### CN neurons exhibit feature selectivity

Next, we examined whether the spatial structure of RFs confer to the firing rate responses of CN neurons a selectivity for specific geometric features, as it does in cortex but not the periphery. To this end, we measured the responses of CN neurons to oriented edges scanned across their RFs. We found that the firing rates of a subset of CN neurons are modulated by orientation, (Figure 6a, Supplementary Figure 2), responding more strongly to edges at some orientations than others. Some neurons are also modulated by direction of movement, responding strongly to a bar scanned in one direction but less so to the same bar scanned in the opposite direction (top right and bottom left neurons in Figure 6a). We quantified the strength of the orientation tuning using a metric – the orientation selectivity index – that takes on a value of 1 when a neuron responds only to a single orientation and 0 when it responds uniformly to all orientations. The degree of orientation selectivity in CN is intermediate between that seen in the nerve – where none exists – and in cortex (Figure 5f). Therefore, the feature selectivity observed in cortex is to some extent inherited from its inputs.

**Figure 6.**
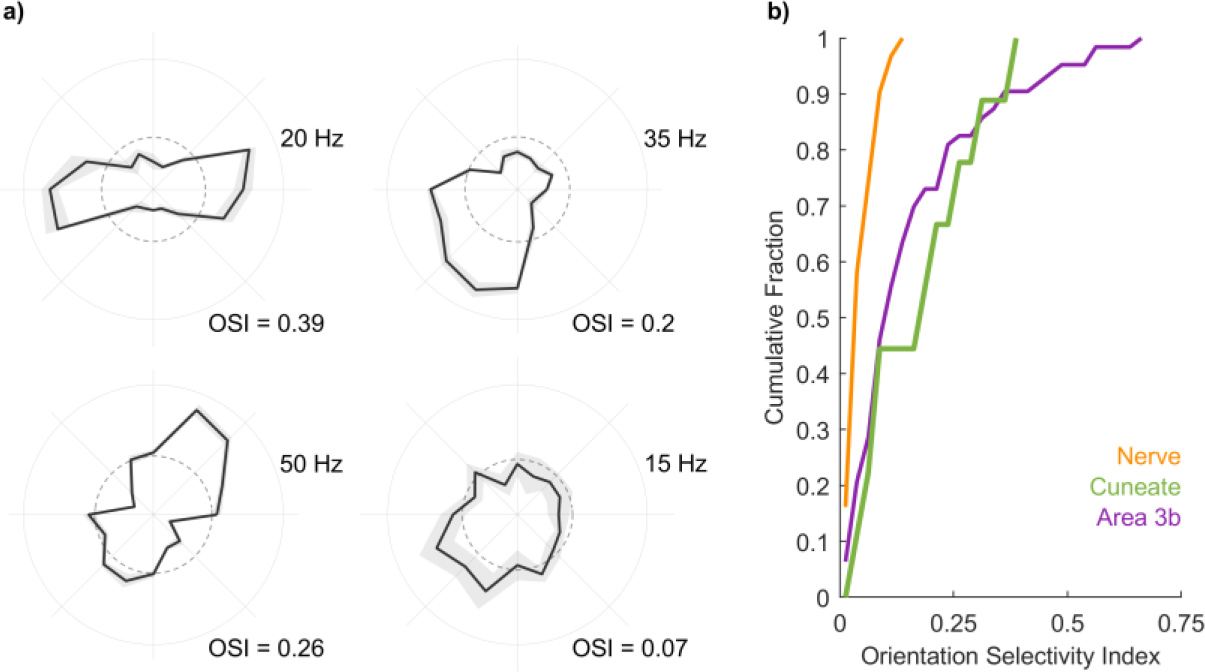
Orientation tuning in CN neurons. a| Response of example CN neurons to oriented edges. The angular coordinate denotes orientation, the radial coordinate denotes firing rate, and the dashed circle denotes the firing rate averaged across conditions. b| Cumulative distribution of orientation selectivity index derived from the responses of nerve fibers, CN neurons, and S1 neurons. CN responses are more strongly tuned for orientation than are nerve fibers but more weakly tuned than are the some S1 neurons.

## Discussion

The objective of the present study was to characterize the tactile representation in CN and to assess the degree to which CN responses reflect computations on their inputs. To these ends, we probed CN responses using stimuli whose representation in the peripheral nerve has been extensively characterized, allowing us to disentangle derived response properties from those inherited from the inputs. Any difference between nerve and CN responses could then be attributed to computations within CN (or possibly to an intervening synapse in the spinal cord (Abraira and Ginty, 2013; Giesler et al., 1984; Liao et al., 2015; Loutit et al., 2021)). We found that CN neurons receive convergent input from multiple tactile submodalities, exhibit spatial and temporal filtering properties that had previously been attributed to cortical processing, and are tuned for behaviorally relevant object features. Comparison of CN responses to their upstream and downstream counterparts suggests that the tactile representation in CN is more similar to its counterpart in cortex than it is to that in the nerve.

### Submodality convergence

Tactile nerve fibers that innervate the glabrous skin of monkeys can be divided into three clearly delineated classes, each with distinct response properties (Delhaye et al., 2018). While each sub-modality might be more responsive to any one stimulus feature, information about most features is distributed over all three sub-modalities and the resulting perceptual experience reflects this integration (Lieber et al., 2017; Muniak et al., 2007; Saal and Bensmaia, 2014; Weber et al., 2013). As might be expected, then, the responses of individual S1 neurons typically reflect convergent input from multiple classes of nerve fibers (Pei et al., 2009). Where this integration might first take place was unclear, however. Studies with cats suggested a lack of submodality convergence in CN (Bystrzycka et al., 1977; Douglas et al., 1978; Ferrington et al., 1987; Gynther et al., 1995; Vickery et al., 1994) whereas studies in rodents conclude that the trigeminal nucleus – a structure analogous to the CN that receives inputs from the face – exhibits submodality convergence at the single cell level (Kaloti et al., 2016; Minnery et al., 2003; Sakurai et al., 2013).

Here, we show that the CN of primates features submodality convergence. Indeed, the majority of CN neurons produce both an SA1-like sustained response to the static component of a skin indentation and an RA/PC-like phasic response at the offset of the indentation. Furthermore, individual CN neurons tend to respond to a wider range of frequencies than do primary afferents of any one class. The submodality convergence observed in somatosensory cortex is thus, at least in part, inherited from its inputs and begins at the earliest processing stage along the dorsal column-medial lemniscus pathway.

### Neural computations

Tactile nerve fibers have small RFs that consist of one or more excitatory hotspots (Johansson, 1978; Vega-Bermudez and Johnson, 1999) and faithfully encode local skin deformations (Saal et al., 2017). In contrast, S1 neurons have larger RFs that comprise excitatory and inhibitory subfields (Bensmaia et al., 2008; DiCarlo et al., 1998; Lieber and Bensmaia, 2019), which confer to them a selectivity for spatial features in their inputs. Individual cortical neurons also act as temporal filters (Saal et al., 2015), which confers to them a selectivity for temporal features in their inputs. The idiosyncratic spatial and temporal filtering properties of individual S1 neurons give rise to a high-dimensional representation of the input in somatosensory cortex, in which different features of grasped objects are simultaneously and unambiguously encoded (Bensmaia et al., 2008; DiCarlo et al., 1998; Lieber and Bensmaia, 2019, 2020; Pei et al., 2010)

Here we show that the spatial and temporal computations observed in cortex are also observed in CN. First, the spatial RFs of CN neurons comprise excitatory and inhibitory subfields and, while somewhat smaller (as expected since CN is upstream from cortex), resemble their cortical counterparts. Second, individual CN neurons process time-varying inputs in a variety of different ways – ranging from integration to differentiation – that are analogous to their cortical counterparts. CN thus contributes to the processing of sensory information and CN neurons exhibit response properties that are qualitatively similar to their counterparts in S1.

### Feature selectivity

The spatiotemporal response properties of S1 neurons confer to them a preference for certain stimulus features. For example, individual S1 neurons exhibit a selectivity for the direction in which objects move across the skin (Gardner and Costanzo, 1980; Pei et al., 2010) or idiosyncratic preferences for different surface textures (Lieber and Bensmaia, 2019). Another well-documented feature selectivity in S1 is for oriented edges: a large proportion of S1 neurons respond preferentially to edges at a specific orientation (Bensmaia et al., 2008). This orientation selectivity is attributed to the neurons’ RF structure, which comprises excitatory and inhibitory subfields, analogous to neurons in primary visual cortex (Hubel and Wiesel, 1962). We show that CN neurons also exhibit orientation selectivity, suggesting that some of the feature selectivity observed in S1 is inherited from its inputs.

Feature extraction results in a sparsening of the stimulus representation, which can result in an overall loss of information (Babadi and Sompolinsky, 2014), unless it is accompanied by an expansion of the size of the neuronal population (Daniel and Whitteridge, 1961). Not surprisingly, the CN is estimated to comprise three to five times more neurons than there are nerve fibers that innervate the corresponding dermatomes, with a preferential expansion of the representation of the hand (Biedenbach, 1972; Corniani and Saal, 2020; Darian-Smith and Ciferri, 2006; Xu and Wall, 1999), also reflected in S1 (Corniani and Saal, 2020), and consistent with observations in other animals (Catania et al., 2011; Lehnert et al., 2021; Wassle et al., 1990). Thus, the expanded neuronal representation in CN is consistent with its role in feature extraction.

### Conclusions

The naïve textbook story is that CN is a simple relay station that does not effect any computations on its inputs but rather transmits them unprocessed. The putative role of CN, if any, has been to provide an opportunity to modulate the gain of the afferent input depending on its behavioral relevance via top down signals (Berkley et al., 1986; Conner et al., 2021). We show that, in addition to this gain modulation, the responses of CN neurons reflect a significant transformation of their afferent inputs, conferring to them properties that were heretofore attributed solely to cortex. CN is thus an active contributor to the process by which ambiguous signals from the periphery are converted into sensory representations that support robust and meaningful percepts and guide behavior.

## Methods

### Neurophysiology

#### Animals & surgical preparation

Neuronal responses were obtained from 7 rhesus macaques (5 males and 2 females, 4-14 years of age, 4-12 kg). Monkeys were anesthetized and placed in a stereotaxic frame with their neck flexed at 90 degrees to provide access to the dorsal brainstem. The foramen magnum was exposed and the inferior aspect of the occipital bone was removed. The dura above obex was resected to reveal the brainstem. All surgical procedures were approved and monitored by the Institutional Animal Care and Use Committee and were consistent with federal guidelines. The neurophysiological methods for the cortical and nerve fiber responses have been previously described (Bensmaia et al., 2008; Harvey et al., 2013; Pei et al., 2009).

#### Neurophysiological recordings

Neuronal activity was monitored using 16-channel linear probes (V-Probe, Plexon, Dallas, TX) and amplified and stored using a Cerebus system (Blackrock Microsystems, Salt Lake City, Utah). Probes were positioned with a stereotaxic system, using the obex as a landmark to locate the CN. Units with receptive fields on the glabrous surface of the hand were isolated. Responses from 143 neurons were obtained across experimental conditions.

### Tactile Stimulation

We presented five classes of stimuli – skin indentations, sinusoidal vibrations, band-pass mechanical noise, scanned random dot patterns, and scanned edges – each with precisely controlled speed, force, frequency, and/or amplitude. In some cases, multiple stimulus classes were delivered while recording from a given neuron. Indentations, sinusoids, and noise stimuli were delivered with a probe (diameter = 1 mm) driven by a custom shaker motor (WESTLING et al., 1976), pre-indented 0.5 mm into the skin. Scanned random dots and edges were presented using a miniaturized version of the drum stimulator (Lieber and Bensmaia, 2019; Weber et al., 2013). Edges were presented using a custom-stimulator that can scan stimuli across the skin in different directions and whose third degree of freedom allows for indentation into and retraction from the skin (see (Pei et al., 2014)). Responses to skin indentation and sinusoids were collected from 4 monkeys, responses to bandpass noise and random dot patterns from 2 monkeys, and responses to edges from 1 monkey.

#### Skin indentations

The amplitude of the ramp and hold indentation was 1 mm and their overall duration was 0.5 seconds, with on- and off- and ramps lasting 25 ms, and separated by a 0.5-s interval. Indentations were presented 100 times.

#### Sinusoids

Sinusoidal vibrations were delivered at 7 frequencies (5-300 Hz) and 10 amplitudes, which spanned the achievable range at each frequency, given the limitations of the stimulator. Each frequency-amplitude combination, lasting 1 s, was presented 5 times in pseudorandom order, separated by a 1-s interstimulus interval, for a total of 350 trials.

#### Bandpass mechanical noise

White Gaussian noise was filtered with different high and low pass frequencies (low: 5-50 Hz, high: 10-200 Hz) to yield 10 unique stimuli (as previously described in (Muniak et al., 2007), each lasting 1 s and separated by a 0.3-s interval.

#### Scanned random dot patterns

Random dot patterns were printed (Form 2, Formlabs, Somerville, MA) on a drum (2.5-in diameter) using previously used geometries and densities (cf. (DiCarlo et al., 1998; Lieber and Bensmaia, 2019)). Patterns were repeatedly scanned across the skin at 80 mm/s. For the first scan, the edge of the pattern was aligned with the estimated the center of the receptive field and indented into the skin by 0.5 mm. For each of 100 subsequent scans, the drum was progressively translated by 0.4 mm along the axis perpendicular to the axis of rotation.

#### Scanned edges

An edge (1-mm high, 1-mm wide, 0.25-mm chamfer) was printed on a miniature drum, whose rotation was driven by a rotational motor. The orientation of the drum on the skin was controlled by a second rotational motor. A third motor controlled the vertical excursion of the drum and allowed for it to be lifted in between changes in orientation. The edge was scanned five times at 80 mm/s at each of 16 orientations (0 – 337.5 degrees with a 22.5 degree spacing).

### Data Analysis

#### Adaptation index

The adaptation index (Pei et al., 2009) indicates the relative firing rate of the sustained and offset periods. SA1 afferents respond to the onset and sustained period while RA and PC afferents respond to the onset and offset transient periods. Thus, the submodality composition of the inputs of a downstream neuron can be measured by taking the ratio of the offset and sustained period. The baseline firing rate was subtracted from both the computed sustained (*fr*_*sus*_) and offset (*fr*_*off*_) firing rates (measured between 0.275-0.375 and 0.505-0.605 seconds respectively). The adaptation index was then computed as:

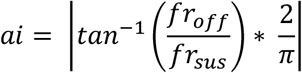

#### Afferent convergence

Given that each class of nerve fibers exhibits a unique frequency response characteristic, we sought to determine if the cuneate responses could be explained by linear combinations of inputs from the three afferent classes. To this end, we simulated the responses of each afferent type to the sinusoidal stimuli used in this study. For this, we used TouchSim, a model that yields millisecond precision reconstructions of the responses of every tactile nerve fiber that innervates the glabrous skin of the hand to arbitrary stimuli delivered to the skin (Saal et al., 2017). The firing rates evoked by each stimulus was then averaged across afferents of the same type. For each cuneate neuron, we used linear regression in the form:

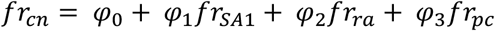

We compared the performance of the full model to that of models that only included one class of nerve fibers. We assessed model performance using 5-fold cross-validation to ensure that models with more parameters did not outperform simpler models due to overfitting. We then compared the best performing single afferent model to that of the full model for each neuron.

To estimate the number of afferents that contribute to the CN response, we performed a regression analysis using the firing rates of individual nerve fibers as regressors. We then used an stepwise linear regression process to determine the optimal combination of inputs. Briefly, on the first iteration, we selected the nerve fiber that had the highest correlation with the CN response. During each subsequent step, we measured the increase in correlation when adding every other afferent in the population, either across classes or within class. We then incorporated the fiber that most improved the regression performance. We proceeded until the addition of an additional regressor failed to improve the model fit more than 5%. To determine the relative contributions of each afferent type to the CN response, we summed the absolute regression coefficient within afferent type and normalized by the summed absolute regression values across afferent types.

#### Spike-triggered average – transfer function

Responses to mechanical noise can be used to estimate the transfer function of a neuron (Saal et al., 2015; Schwartz et al., 2006). Accordingly, we simulated the responses of all the nerve fibers that innervate the glabrous skin of the hand to the bandpass mechanical noise used in the neurophysiological experiments and averaged their responses across fibers of each class. We then performed a spike triggered average (STA) (Schwartz et al., 2006) of the response of each afferent conditioned on each spike in CN. That is, the response of each afferent population over the 100 ms preceding each CN spike was averaged across CN spikes. The resulting filter was smoothed using a Gaussian filter (sd = 5 ms) and the baseline firing rate was subtracted. We then standardized the resultant filter for each cuneate-afferent pair with respect to the period between 100 and 50 ms before the cuneate spike, which we expect to reflect noise. We then thresholded (Z > 2) the Z-scored probabilities and computed the magnitude and width of the filters.

#### Harmonic ratio

To determine the extent to which CN response to sinusoids were phase-locked, we computed the harmonic ratio of the response to each stimulus. Excluding responses with fewer than 5 spikes, we performed a Fast Fourier transform (FFT) of the peristimulus spike histogram (binned at 1/5f;), averaged the mean amplitude at the fundamental frequency (*A*_*f*_) and its first harmonic (*A*_*h1*_), and divided the resulting value by the median amplitude across all frequencies (*Ã*):

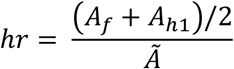

where:

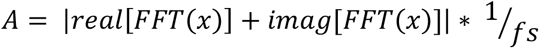

We repeated this analysis for Poisson spike trains to obtained a distribution of harmonic ratios obtained by chance.

#### Spatial receptive fields

We used standard techniques to estimate the spatial receptive field of each cuneate neuron (cf. (DiCarlo et al., 1998; Lieber and Bensmaia, 2019)). In brief, we averaged the 16 mm x 16 mm swath of the random dot pattern that impinged on the skin at the time of each cuneate spike. To remove the curvature of the drum reflected in the resulting STA, we subtracted a 2^nd^ order polynomial plane from it. The resultant STA was then standardized and thresholded to isolate excitatory and inhibitory lobes.

To identify the number of subfields for cortical and cuneate receptive fields, we fitted 2D Gaussians to each RF. Each Gaussian subfield had the following form:

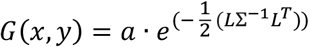

where *L* = [*x* – *μ*_*x*_, *y* – *μ*_*y*_], Σ = CovMat(*σ*_*x*_, *σ*_*y*_, θ), *a* is the amplitude (*a* > 0 denotes an excitatory patch, *a* < 0 an inhibitory one), (x, y) denote the medial-lateral and proximal-distal locations on the skin surface, respectively, (*μ*_*x*_, *μ*_*y*_) represent the center of the Gaussian, (*σ*_*x*_, *σ*_*y*_) its standard deviations along the two axes, and *θ* its orientation.

Therefore, every RF is described by a total of *N* × 6 parameters (6 parameters for each Gaussian component: *a, μ_x_, μ_y_, σ_x_, σ_y_*, and *θ*; and *N* Gaussian components). Nonlinear least-squares optimization was used to find the best parameters. *N* represents the minimum number of Gaussian subfields needed to achieve *R*^*2*^ > 0.9 of 90% of the maximum achievable *R*^*2*^.

#### Orientation tuning

Spiking responses evoked at each orientation were aligned to a reference stimulus trace consisting of 6 Gaussians spaced according to the stimulus speed. The spike rate evoked by the stimulus, centered around the peak response, was averaged over a window of 314 ms, corresponding to 6.2 mm of travel (5% of a complete rotation of the drum), though the window size did not affect the results over a wide range (Supplementary Figure 2). The tuning of each neuron was gauged using an Orientation Selectivity Index, given by:

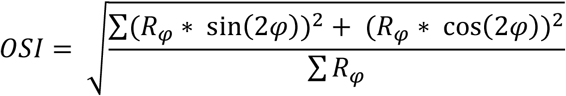

Where ϕ is the orientation of the stimulus and *R*_*φ*_ is the firing rate at that orientation. Reliability of the OSI was tested using a permutation test, for which neural responses were shuffled 10000 times and the OSI recomputed.

## Acknowledgments

We would like to thank Amit Ayer for help with the surgical procedures. This work was supported by NINDS grants NS122333 (SJB), NS095162 (SJB, JMR, LEM) and NS096952 (AKS).

## Supplemental Figures

**Supplementary Figure 1:**
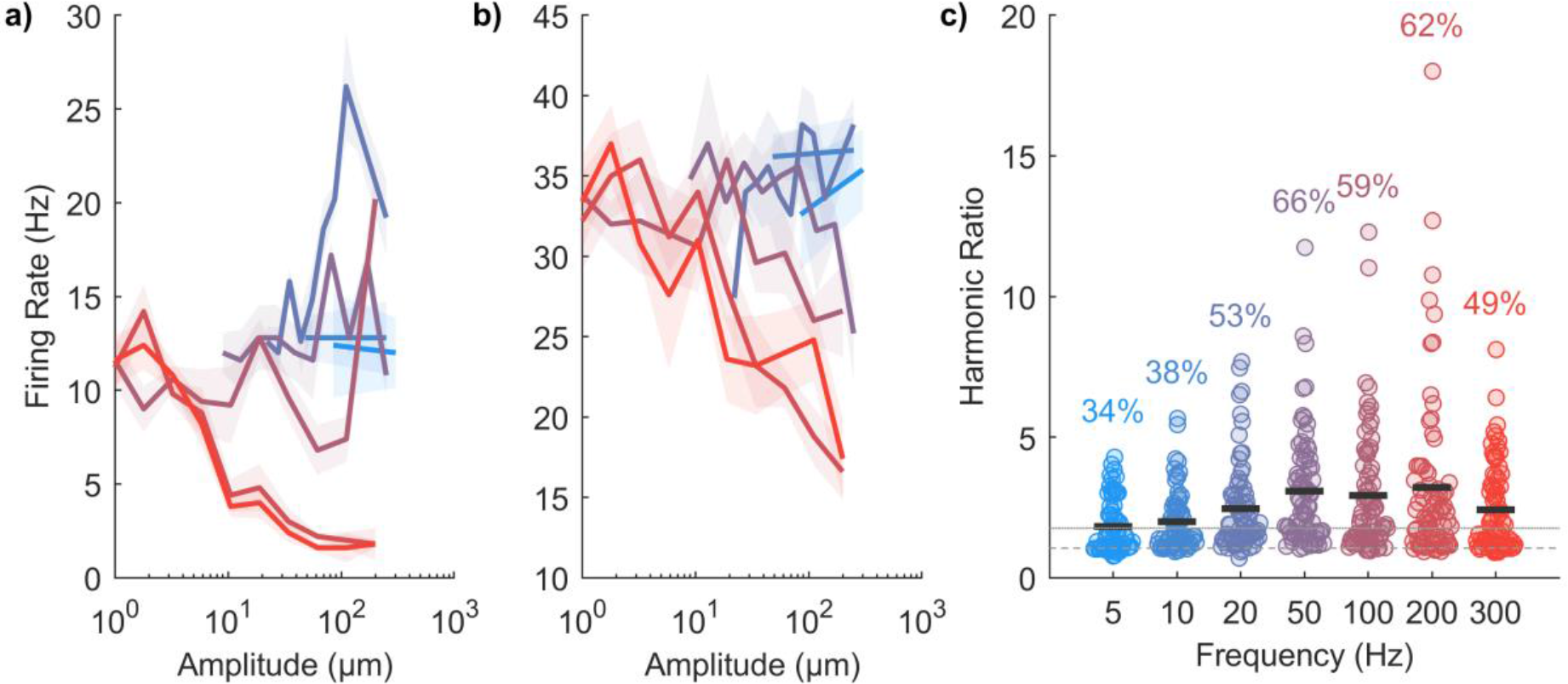
Frequency characteristics of CN neurons. a & b| Example rate-intensity plots for two CN neurons. The neuron in panel *a* is sometimes excited, sometimes inhibited by skin vibrations depending on their frequency while the neuron in panel *b* exhibits only suppression. c| The proportion of neurons that are significantly tuned to each frequency. The dashed line indicates the mean harmonic ratio of a Poisson neuron (∼1), and the dotted line above is the value 3 standard deviations above the mean.

**Supplementary Figure 2:**
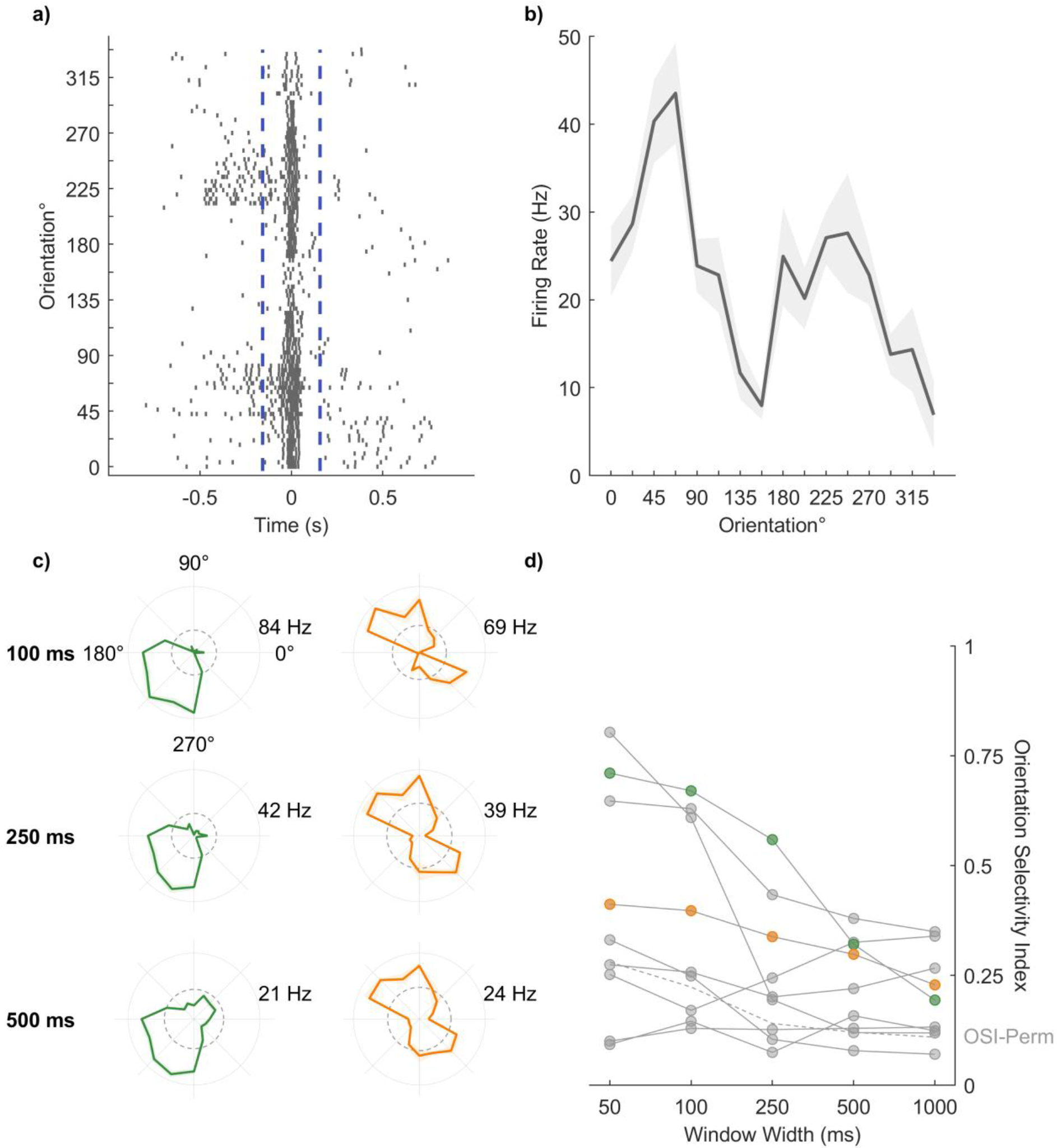
Orientation tuning is stable with respect to window size. a| Raster plot of the responses of an example CN neuron to edges at each orientation aligned to the peak response. b| Tuning curve for the neuron shown in panel a. c| Polar plots for 2 example neurons computed from the responses averaged over 3 time windows. d| Vector strength for all neurons computed over different windows. Example neurons from panel *c* are highlighted in the corresponding color. OSI-Perm indicates the orientation index averaged across neurons when the firing rates are shuffled across trials.

